# Chlorzoxazone, A BKCa Channel Agonist, Rescues The Pathological Phenotypes Of Williams-Beuren Syndrome In A Preclinical Model

**DOI:** 10.1101/2020.12.22.423977

**Authors:** Marion Piquemal, Noura Abdulkarim-Abdalla, Paula Ortiz-Romero, Valerie Lemaire-Mayo, Wim E. Crusio, Eric Louette, Victoria Campuzano, Susanna Pietropaolo

## Abstract

Williams-Beuren syndrome (WBS) is a rare developmental disorder caused by the deletion of a 1.5 Mb region in chromosome 7 (7q11.23). WBS has been recently modelled by a mutant mouse line having a complete deletion (CD) of the equivalent locus on mouse chromosome 5, thus resembling the genetic defect found in WBS patients. CD mice have been shown to have physical and neurobehavioral abnormalities that recapitulate most of the symptoms associated with human WBS, including cardiovascular, motor, social, emotional and sensory alterations. This model has been largely used to investigate the etiopathological mechanisms of WBS; nonetheless, pharmacological therapies for this syndrome have not been identified yet. Here we propose a novel treatment for WBS, chlorzoxazone (CHLOR), i.e., a molecule targeting calcium-activated large conductance potassium (BKCa) channels, since a reduction in the expression of these channels has been recently described in neurons from WBS patients, as well as in other rare developmental pathologies. Our results demonstrate both the acute and chronic effects of CHLOR on some major pathological phenotypes of CD mice, including several behavioural alterations and cardiac hypertrophy. We conclude that BKCa channels are a therapeutic target of high potential for clinical applications and are likely to play a key role in the etiopathology of WBS.

## INTRODUCTION

Williams-Beuren syndrome (WBS) is a rare developmental disorder, caused by the hemizygous deletion of 26–28 contiguous genes on chromosome band 7q11.23 (Schubert 2009). WBS patients present specific physical abnormalities, including craniofacial alterations and growth retardation, together with cardiovascular anomalies, including a generalized arteriopathy with aortic stenosis and hypertension, and cardiac hypertrophy (Pober 2010). Patients also show elevated anxiety levels, acoustic and sensory alterations, hypotonia and motor problems, and a characteristic cognitive profile including hypersociability and deficient visuospatial abilities (Martens, Wilson et al. 2008). Recently, a mouse model recapitulating the complete chromosomic deletion (CD mouse) and most of the cardio-vascular and neurobehavioural phenotypes of WBS has been engineered (Segura-Puimedon, Sahun et al. 2014). This pre-clinical model has largely contributed to advance our understanding of the molecular pathological mechanisms involved in WBS (Borralleras, Sahun et al. 2015, Borralleras, Mato et al. 2016, Ortiz-Romero, Borralleras et al. 2018, Dasilva, Navarro-Guzman et al. 2020, Jimenez-Altayo, Ortiz-Romero et al. 2020), including reduced brain development, altered synaptic mechanisms and morphological abnormalities in specific brain areas. The CD model has also been recently employed to test the efficacy of potential novel therapeutic approaches to WBS, especially focusing on non-pharmacological interventions [e.g., based on natural polyphenolic extracts, (Ortiz-Romero, Borralleras et al. 2018)]. Nonetheless, effective pharmacological treatments acting on both the behavioural and the cardiovascular phenotypes of WBS patients are still lacking, thus highlighting an urgent need of identifying novel powerful therapeutic targets.

Potassium (K+) channels are of high interest for the functionality of the nervous system, as they play a crucial role in regulating the triggering of neurons and the release of neurotransmitters (Latorre and Brauchi 2006). This modulation of synaptic transmission and electrical discharge at the cellular level is due in particular to the expression of large-conductance Ca2+-activated K+ (BKCa) channels (Sah and Faber 2002). The opening of these channels causes a movement towards the potassium equilibrium potential and thus accelerates the repolarization of the action potentials, allowing faster stimulation (Hu, Shao et al. 2001). In addition, BKCa channels modulate the activity of dendrites as well as astrocytes and microglia. Beside their role in the central nervous system, BKCa channels are also involved in smooth muscle contraction and endocrine cell secretion (Jackson 2005, Tykocki, Boerman et al. 2017). These channels are assembled as hetero-octamers of four α-subunits (KCNMA1 protein) and four auxiliary regulatory β-subunits, and are activated by membrane depolarization and increased intracellular Ca2+ concentration (Gribkoff, Starrett et al. 2001). Converging lines of evidence support the role of BKCa channels in the etiopathology of several neurodevelopmental disorders (NDDs)(Lee and Jan 2012). A KCNMA1 gene disruption was described in a subject with autism, associated with a functional defect of BKCa channels (Laumonnier, Roger et al. 2006). We also demonstrated a reduced expression of the KCNMA1 protein in another rare NDD, i.e., Fragile X syndrome (FXS), both in patients and in the related Fmr1-KO mouse model (Hebert, Pietropaolo et al. 2014), together with a reduced functionality of brain BKCa channels (Zhang, Bonnan et al. 2014). Recently, reduced expression of KCNMA1 has been reported in neurons from WBS patients (Khattak, Brimble et al. 2015). These data strongly support the relevance of BKCa channels as a potential therapeutic target common to multiple NDDs (N’Gouemo 2014), suggesting that these disorders should be considered as “channelopathies” (Bailey, Moldenhauer et al. 2019). Preclinical studies have indeed demonstrated that the administration of molecules acting as BKCa channel openers may rescue the brain and behavioural abnormalities of Fmr1-KO mice (Hebert, Pietropaolo et al. 2014, Zhang, Bonnan et al. 2014, Carreno-Munoz, Martins et al. 2018). Nonetheless, the relevance of BKCa channels as therapeutic targets for WBS has never been specifically addressed so far.

Hence, here we evaluated whether the administration of one molecule acting on BKCa channels could represent an effective pharmacological strategy to treat WBS symptoms, including behavioural, physical, brain and cardiovascular alterations. To this end we focused on Chlorzoxazone, (5-chloro-2,3-dihydro-1,3-benzoxazol-2-one; CHLOR), as we have recently demonstrated the efficacy of this agonist of BKCa channels (Liu, Lo et al. 2003) in the Fmr1-KO mouse model of FXS (Lemaire-Mayo, Piquemal et al. 2020). CHLOR is a classical drug, approved from FDA as a muscle relaxant (Hohmann, Blank et al. 2019), acting on the spinal cord and subcortical areas of the brain (Martindale 1993). Hence, the main advantage of CHLOR is that it could be rapidly employed in clinical trials through a drug repurposing strategy. In fact, recent pre-clinical studies have highlighted the emerging potential role of CHLOR in healthcare, demonstrating its immunomodulatory (Deng, Li et al. 2020) and neuroprotective properties (Bai and Ma 2020). The present study provides with the first characterization of the therapeutic effects of CHLOR administration in the CD mouse model of WBS. To this end, we assessed both acute and chronic effects on the WBS-like behavioural alterations of CD mice, including motor, social, emotional and sensory deficits. Furthermore, we evaluated the effects of chronic treatment with CHLOR on the physical, brain and cardiovascular abnormalities of CD mice.

## METHODS

### Animals

Subjects were adult (4-5 months old) CD mutant heterozygous mice and their wild-type littermates (maintained on B6 background). Males were more extensively tested, as they are preferentially employed in studies on mouse models of NDDs, including our previous work with CHLOR on the Fmr1-KO model of FXS (Lemaire-Mayo, Piquemal et al. 2020). Nonetheless, since WBS affects individuals of both sexes, we replicated part of the experiments in female mice, as described in the Supplementary material section.

Mice tested in studies 1 and 2 were bred in our animal facility of Bordeaux University, from CD breeders, heterozygous for the CD mutation (Segura-Puimedon, Sahun et al. 2014). Breeding trios were formed by mating two heterozygous CD males with a wild-type C57BL/6J female purchased from Janvier (Le Genest St Isle, France). After 2 weeks the sire was removed and the females were single caged and left undisturbed until weaning of the pups. Mice were weaned at 21 days of age and group-housed with their same-sex littermates (3–5 animals/cage). On the same day, tail samples were collected for DNA extraction and subsequent PCR assessment of the genotypes as previously described (Segura-Puimedon, Sahun et al. 2014). Only litters including males of both genotypes (WT and CD) were used for experiments.

Mice used in study 3 were bred and maintained in the animal facility at the University of Barcelona; CD heterozygous mice, obtained as previously described (Segura-Puimedon, Sahun et al. 2014), were crossed with Thy1-YFP transgenic mice (B6.Cg-Tg(Thy1-YFPH)2Jrs/J, Jackson Laboratory) (Feng, Mellor et al. 2000) in order to label pyramidal neurons and allow direct evaluation of dendritic length.

CD1 female mice (12±1 weeks old) purchased from Janvier (Le Genest St Isle, France) were used as social stimuli during the social interaction test. This strain has been selected since it was previously largely employed as social stimuli in several social studies from our group, including those on Fmr1-KO mice (Pietropaolo, Guilleminot et al. 2011, Pietropaolo, Goubran et al. 2014, Oddi, Subashi et al. 2015), because of its high sociability levels (Moles and D’Amato F 2000). Mice were group-housed (4-5 animals/ cage) and left undisturbed upon arrival at least one week before the social interaction test.

All animals were housed in polycarbonate standard cages (33×15×14 cm in size; Tecniplast, Limonest, France), provided with litter (SAFE, Augy, France) and a stainless steel wired lid, and enriched with a cotton nestlet. Food (SAFE, Augy, France) and water were provided ad libitum. The animals were maintained in a temperature (22°C) and humidity (55%) controlled vivarium, under a 12:12 hr light–dark cycle (lights on at 7 a.m.). All experimental procedures were in accordance with the European Communities Council Directive of 24 November 1986 (86/609/EEC) and local French and Spanish legislation.

### Behavioural tests

To evaluate the therapeutic potential of CHLOR we have chosen behavioural tests that have revealed altered phenotypes in CD mice based on previous data from Dr. Campuzano’s team and from our pilot studies. These phenotypes include motor coordination deficits (rotarod test), hypo-activity (the open field test), hyper-sociability (the direct social interaction test for social habituation), altered natural occurring behaviours (test for nesting abilities in a novel environment and marble burying test), increased anxiety (self-grooming in a novel environment) and alterations in sensory responsiveness (acoustic startle test). The choice of these tests was based on their ability to reveal WBS-like phenotypes in CD mice, and to be sensitive to the therapeutic effects of the molecule CHLOR in a mouse model of another neurodevelopmental disorder, i.e., the Fmr1-KO model for FXS (Lemaire-Mayo, Piquemal et al. 2020). Furthermore, these tests were selected since they allow a rapid assessment of mouse behavior, with minimal manipulation of the subjects, thus allowing quick and repeated testing of the same batch of mice, concomitantly reducing the number of subjects needed.

All behavioral tests (except nest building assessment) were carried out during the light phase of the cycle. Mice were habituated to the experimental room prior to all behavioral tests, being individually housed in standard polycarbonate cages provided with sawdust, food, and water bottles and left undisturbed for at least 10-15 min before testing began.

#### Rotarod test

An electrical accelerating rotarod for mice (Model 7650; Ugo Basile, Stoelting, Wood Dale, IL) was used to evaluate motor coordination. Mice were placed on the rotating drum at the baseline speed of 4 rpm. During the 5-min observation period, the speed of rotation increased linearly to 40 rpm. Mice were given three trials, with an intertrial interval of one hour. Each trial ended when the mouse fell from the apparatus or when 5 min had elapsed.

The latency to fall from the rotating drum was recorded, averaged across trials and ln-transformed to better conform to the assumptions of parametric ANOVA..

#### Open field test

This test is used to assess locomotor activity and exploration in laboratory mice (Belzung, Leman et al. 2005). The apparatus consisted of 4 white opaque plastic arenas (42×26×15cm). Each mouse was placed in the center of the arena and left free to explore it for 5 minutes. Automated Tracking of the videos obtained from a camera above the open field was performed with Ethovision (version 11, Noldus Technology, Wageningen, Netherlands) to analyze the total distance travelled.

#### Direct social interaction test and self-grooming assessment

Social interaction was assessed in a 30×15×14 cm cage, covered by a flat metal grid and with approximately 3 cm of sawdust on the floor, where testing subjects were previously isolated for one hour. During the first 10 min of isolation in the novel cage, behaviour of the subject was videorecorded and the time spent performing self-grooming was afterwards assessed with Observer XT (version 7, Noldus, The Netherlands).

An unfamiliar CD1 stimulus female (3 months-old), was then introduced into the testing cage and left there for 6 min. Stimulus females were housed in unisexual groups in a female-only animal room and were in the non-estrous phase when tested (as assessed by the analysis of vaginal smears on testing day).

Testing sessions were recorded and videos analyzed with Observer XT (version 7, Noldus, The Netherlands). Affiliative behaviors only of the testing subject were evaluated, including sniffing the head and the snout of the partner, its anogenital region, or any other part of the body; allogrooming (grooming the partner) and traversing the partner’s body by crawling over/under from one side to the other. Social habituation was assessed through the analysis of the time spent in affiliative behaviours across 2-min bins.

#### Nest-building behaviour in a novel environment

The apparatus consisted of an unfamiliar Plexiglas cage (30×15×14 cm), covered by a flat metal grid, with approximately 3 cm of sawdust on the floor, provided with food and a water bottle. Mice were singly placed in the apparatus in the presence of nesting material (one Nestlet, 2.7 g, 2.5 × 2.5 cm and 5mm thick compressed cotton; identical to the material provided in the home cage) and left undisturbed overnight. Nest building scoring was performed by a trained experimenter blinded to the genotype and treatment of the animals, using the following standardized scoring scale (Halene, Ehrlichman et al. 2009, Mo, Renoir et al. 2015, Carreno-Munoz, Martins et al. 2018): 1: Nestlet not noticeably shredded; 2: Nestlet 10 to 50% shredded, not used as a nest; 3: Nestlet shredded 50 to 90%, but the shredded material remains scattered in the cage and is not used as a nest; 4: Nestlet shredded >90%, and shredded material used as a flat nest; and 5: Nestlet shredded >90% and used as a rounded nest with sides covering the mouse.

#### Marble burying

The test was conducted in a polycarbonate rat cage filled with bedding to a depth of 5cm and lightly tamped down. A regular pattern of 20 glass marbles (five rows of four each) was placed on the surface of the bedding prior to each test. An animal was individually placed in the testing cage for 20 min and the number of buried marbles (>2/3 marble covered) was counted.

#### Acoustic startle test

The apparatus consisted of four acoustic startle chambers for mice (SR-LAB, San Diego Instruments, San Diego, CA, USA). Each comprised a nonrestrictive cylindrical enclosure made of clear Plexiglas attached horizontally on a mobile platform, which was in turn resting on a solid base inside a sound-attenuated isolation cubicle. A high-frequency loudspeaker mounted directly above the animal enclosure inside each cubicle produced a continuous background noise of 65/ 66 dBA and various acoustic stimuli in the form of white noise. Vibrations of the Plexiglas enclosure caused by the whole-body startle response of the animal were converted into analog signals by a piezoelectric unit attached to the platform. These signals were digitized and stored by a computer. The sensitivity of the stabilimeter was routinely calibrated to ensure consistency between chambers and across sessions. A session began when the animals were placed into the Plexiglas enclosure (to which they were habituated for 5 min in the absence of acoustic stimuli the day before the test, in order to minimize the stress).

We used two protocols to evaluate acoustic startle test in our CD mice. First, we performed the test with a protocol involving low intensity stimuli that has been used so far only in Fmr1-KO mice where it has allowed detecting the sensory-hyper responsiveness of these FX mice (Michalon, Bruns et al. 2014, Zhang, Bonnan et al. 2014, Gaudissard, Ginger et al. 2017). After 5 min of habituation, mice were presented with pulses of white sound of 20 ms duration and varying intensity: +6, +12 +18 and +24 dB over the 66db background level (namely 72, 78, 84 and 90 dB). Each intensity was presented 8 times, in a randomized order with variable intervals (10 sec to 20 sec) between the onset of each pulse. As the results obtained with this first procedure did not show a convincing phenotype of our male mutants, we then apply another protocol providing an extensive assessment of acoustic startle response from low (69 dB) to high (120 dB) intensities. Ten pulse intensities were used (over a background noise of 65dB): 69, 73, 77, 81, 85, 90, 95, 100, 110 and 120dB lasting for either 20 or 40 ms. Mice were acclimatised to the apparatus for two minutes before the first trial began. The first six trials consisted of six trials at 120 dBA, with three trials at each of the two possible stimulus durations. These trials served to stabilize the animals’ startle response, and were not included in the overall analysis. Subsequently, the animals were presented with 5 blocks of discrete test trials. Each block consisted of twenty pulse-alone trials, one for each intensity and duration, presented in a pseudorandom order. The inter-trials interval was variable (9-19 s) with an average duration of 14 s.

For all procedures, a total of 130 readings of the whole-body startle response were taken at 0.5-ms intervals (i.e., spanning across 65 ms), starting at the onset of the pulse stimulus. The average amplitude (in mV) over the 65 ms was used to determine the stimulus reactivity and further averaged across trials. A natural logarithmic transformation was applied in order to fulfill the normality criteria requested by parametric ANOVA.

### Drug preparation

All injectable solutions were freshly prepared on each experimental day. CHLOR (Sigma Aldrich, France) was dissolved in saline solution containing 1.25% DMSO (Sigma Aldrich, France) and 1.25% Tween80 (Sigma Aldrich, France). The same solution without drugs was used for the vehicle (VEH) control group. CHLOR was administered at the dose of 5mg/Kg in both acute and chronic studies, based on our previous dose-response studies in WT and Fmr1-KO mice (Lemaire-Mayo, Piquemal et al. 2020).

### Brain and heart analyses

After cardiac perfusion with 1x PBS followed by 4% paraformaldehyde, brains and hearts were removed. Brains were postfixed in 4% paraformaldehyde for 24 hours at 4ºC, in PBS for 24 hours at 4ºC, and then crioprotected in 30% sucrose for 24 hours at 4ºC. Serial coronal brain sections (150μm) were collected on a glass slide and directly mounted with Mowiol. Hearts were postfixed in 4% paraformaldehyde for 48 hours and then paraffin embedded. Serial coronal heart sections (8μm) were collected on a glass slide and stained with a regular hematoxylin-eosin protocol.

For morphological analysis 1360×1024 images of the cortex, and CA1 hippocampus were obtained with an Olympus DP71 camera attached to an Olympus BX51 microscopy with an Olympus U-RFL-T source of fluorescence at −10x magnification. Measures from six sections per animal were averaged. Images of hematoxylin-eosin stained heart samples were obtained using visible light with an Olympus DP71 camera attached to an Olympus MVX10 MacroView Upright Microscope (zoom factor 1,25) Cardiomyocytes’ areas were measured using ImageJ software.

### Statistical analysis

All data were analyzed with ANOVA using treatment and genotype as the between-subject factors and adding trials (for the rotarod), stimulus intensity (acoustic startle), 2-min bins (social interaction) as the within-subject factors. Post-hoc comparisons were performed when a significant interaction genotype x treatment was found using Tukey’s-Kramer test. Otherwise, separate one-way ANOVAs in each treatment group with genotype as the between subject factor were conducted, if appropriate. All analyses were carried out using Statview and PASW Statistics 18. All data are expressed as mean± SEM; p was set at 0.05.

## RESULTS

### Study 1: Behavioural effects of acute CHLOR treatment in CD mice

Mice received a single i.p. injection of VEH solution or CHLOR (5mg/Kg). One hour after injection, animals were submitted to the rotarod test, the open field, the social interaction test, and acoustic startle test with low intensity stimuli (72-90 dB).

As expected, CD mice treated with VEH showed motor alterations, evident as an overall reduction in the latency to fall from the rotarod and hypoactivity in the open field (Fig.1-A and -B), and deficits in social habituation (Fig.1-C). The abnormalities in motor coordination in the rotarod test (Fig.1-A) were attenuated by acute CHLOR administration that was able to significantly improve the motor learning abilities of mutant mice on the 3^rd^ trial [interaction genotype x treatment: F(1,41)=4.84, p<0.05 and treatment x trial F(2,84)=3.32, p<0.05; genotype x treatment effect in separate ANOVA on trial 3: F(1,43)=19.34, p<0.0001; post-hoc: CD-VEH versus WT-VEH and versus CD-CHLOR]. Acute CHLOR fully eliminated the hypoactivity of CD mice in the open field [Fig.1-B; interaction genotype x treatment: F(1,45)=6.31, p<0.01; post-hoc: CD-VEH versus WT-VEH and versus CD-CHLOR]. CHLOR also restored social habituation in the direct social interaction test (Fig.1-C), with CHLOR-treated CD mice showing a time-dependent reduction in social investigation similar to WT mice [genotype x treatment x 2-min bins: F(2,42)=3.33, p<0.05; post-hoc: CD-VEH versus WT-VEH and versus CD-CHLOR on the last time bin]. CD mice treated with VEH showed a tendency to reduce sensory responsiveness in the acoustic startle test, especially at the highest intensities (Fig.1-D). Nonetheless, this phenotypic difference did not reach statistical significance [genotype x stimulus intensity: F(3,138)=2.29, p=0.08; treatment effect and its interactions: n.s.]; hence, we employed a more extensive evaluation of acoustic startle in our subsequent study with chronic administration.

**Fig. 1:**
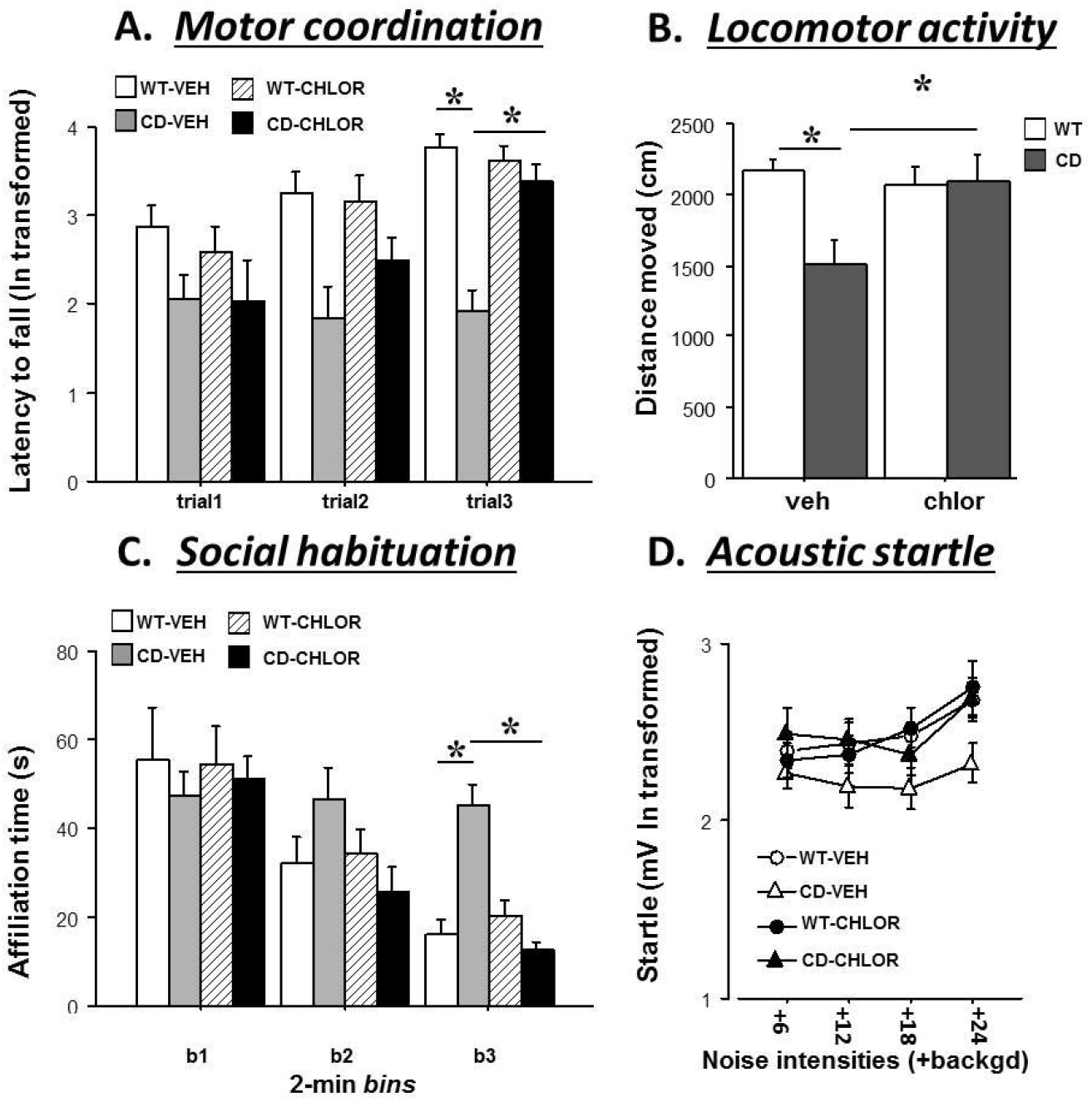
Acute behavioural effects of chlorzoxazone (CHLOR; 5mg/Kg): Behavioural tests were conducted 1h after a single i.p. injection and they included rotarod test for motor coordination (A), open field test for locomotor activity (B), social habituation during direct social interaction with a WT female (C), and acoustic startle response test (D) with low intensity stimuli (background level=66dB). n=6-14; *=p<0.05. Data are expressed as Mean±SEM.

### Study 2: Behavioural effects of chronic CHLOR administration in CD mice

Mice received a daily i.p. injection of VEH solution or CHLOR (5mg/Kg) for 10 days before the beginning of behavioural testing. This experimental protocol was previously used in our study in Fmr1-KO mice where it showed marked behavioural effects of CHLOR (Lemaire-Mayo, Piquemal et al. 2020). Behavioural tests were carried out 24hs after the last injection, in order to avoid acute confounding effects. They included rotarod test, assessment of overnight nest building, evaluation of self-grooming in a novel environment, direct social interaction and acoustic startle (69-120 dB) tests. Nest building behavior is an early and highly sensitive indicator of behavioural dysfunction (Gaskill, Karas et al. 2013, Jirkof 2014, Kraeuter, Guest et al. 2019, Neely, Pedemonte et al. 2019), altered in several models of neuropsychiatric disorders (Halene, Ehrlichman et al. 2009, Mo, Renoir et al. 2015, Carreno-Munoz, Martins et al. 2018). Self-grooming is also a naturally occurring behavior, that is more markedly expressed as an emotional response to a novel environment; it is therefore used as a sensitive indicator of enhanced anxiety, especially in genetic mouse models where other non-ethologically based tests fail to reveal emotional phenotypes [e.g., the Fmr1-KO mouse model for FXS, (Gantois, Khoutorsky et al. 2017, Lemaire-Mayo, Piquemal et al. 2020)].

As expected, CD mice treated with VEH showed deficits in motor abilities and daily activities (Fig.2-A and B), enhanced anxiety levels (Fig.2-C), reduced social habituation (Fig.2-D) and altered acoustic startle response (Fig.2-E); all these abnormalities were attenuated or fully rescued by chronic CHLOR administration. The abnormalities of CD mice in motor coordination in the rotarod test (Fig.2-A) were attenuated by chronic CHLOR administration, improving motor learning on the 3^rd^ trial [interaction genotype x treatment x trial: F(2,66)=2.95, p=0.06; genotype x treatment effect in separate ANOVA on trial 3: F(1,33)=9.42, p<0.01; post-hoc: CD-VEH versus WT-VEH and versus CD-CHLOR]. CHLOR eliminated the reduced nesting abilities of CD mice [Fig.2-B; interaction genotype x treatment effects: F(1,34)=4.16, p<0.05; post-hoc: CD-VEH versus WT-VEH and versus CD-CHLOR], as well as their enhanced anxiety, i.e., self-grooming levels in a novel environment [Fig.2-C; interaction genotype x treatment effects: F(1,33)=7.56, p<0.01; post-hoc: CD-VEH versus WT-VEH and versus CD-CHLOR]. CHLOR also restored social habituation in the direct social interaction test (Fig.2-D), as CD mice treated with CHLOR showed a time-dependent reduction in social investigation similar to WT mice, an habituation process that was absent in CD-VEH animals [genotype x treatment x 2-min bins: F(2,68)=5.41, p<0.01; post-hoc: CD-VEH versus WT-VEH and versus CD-CHLOR on the last time bin]. Finally, CHLOR rescued the acoustic hypo-responsiveness of CD mice that was mostly evident at the highest stimulus intensities [Fig.2-E; genotype x treatment x stimulus intensity: F(9,306)=5.75, p<0.0001; post-hoc: CD-VEH versus WT-VEH and versus CD-CHLOR on the two highest intensities].

**Fig. 2:**
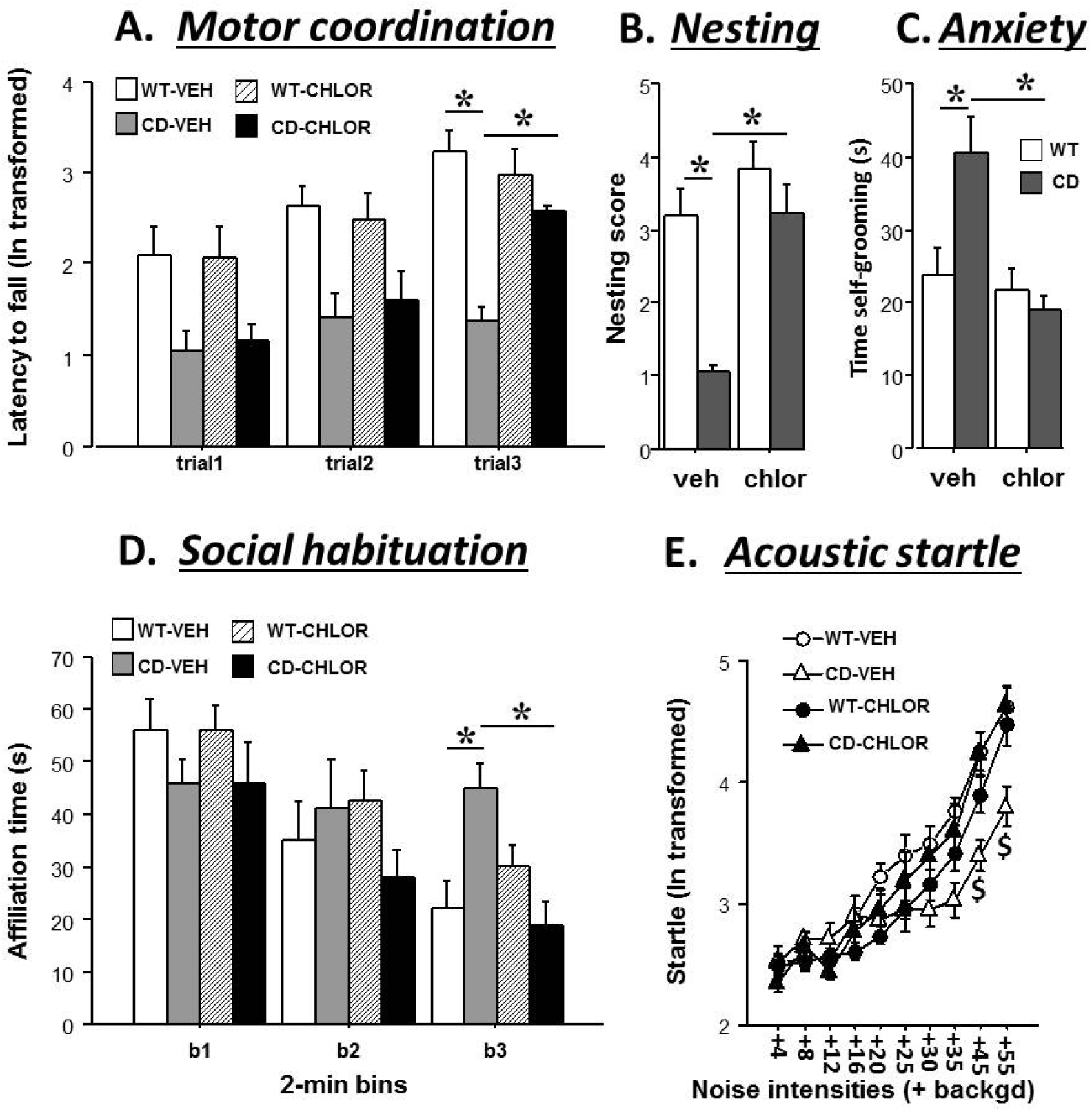
Chronic behavioural effects of chlorzoxazone (CHLOR; 5mg/Kg) in male mice: Behavioural tests were conducted after 10 days of i.p. injections, with a 24hs interval from the last administration. They included the rotarod test for motor coordination (A), nest building assessment (B), anxiety levels measured by levels of self-grooming in a novel environment (C), social habituation during direct social interaction with a WT female (C), and acoustic startle response test (D, background level: 65dB). n=8-10; *=p<0.05; $= versus CD-CHLOR and WT-VEH. Data are expressed as Mean±SEM.

**Fig. 3:**
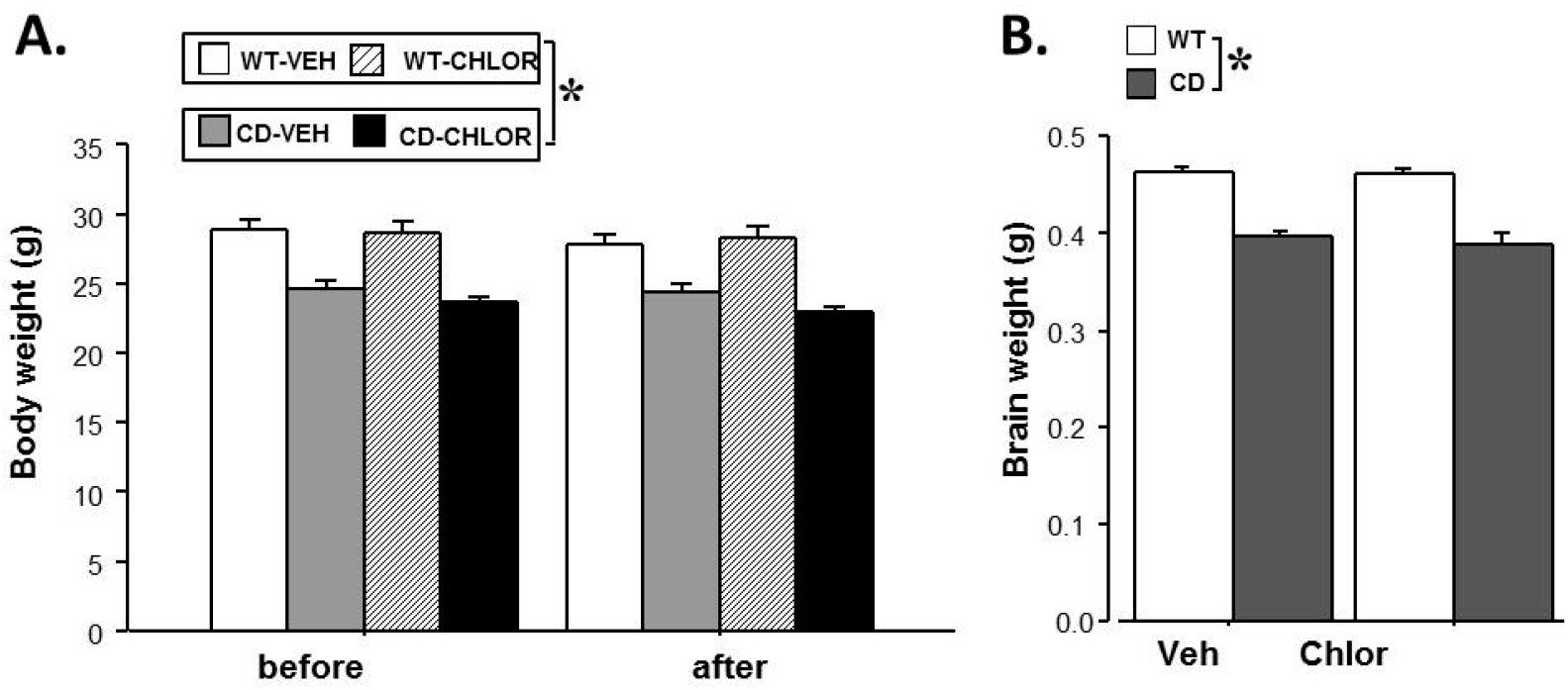
Chronic effects of chlorzoxazone (CHLOR; 5mg/Kg) on body and brain weight in male mice: Final measures of body weight (A) and brain weight (B) were taken after 10 days of i.p. injections, with a 24hs interval from the last administration, n=5-6; *=p<0.05. Data are expressed as Mean±SEM.

### Study 3: Chronic effects of CHLOR on body, heart and brain abnormalities

Mice received a daily i.p. injection of CHLOR during 10 days as in study 2 before being sacrificed for heart and brain morphological assessment. Before sacrifice, body weight was evaluated and mice underwent the marble-burying test, to confirm the behavioural chronic effects of CHLOR on this batch of mice. This test assesses a natural occurring defensive behaviour of laboratory mice, consisting in covering threatening objects, though it is sometimes employed as a test for anxiety or repetitive/obsessive behaviours. Therapeutic effects of CHLOR on the behaviour of CD mice were confirmed in this test, as CD-VEH animals displayed lower levels of marble burying, and these were normalized by CHLOR chronic treatment [interaction genotype x treatment: F(1,17)= 34.96, p<0.0001, Mean±SEM of number of buried marbles were: WT-VEH=17.2±0.74, CD-VEH=4.2±1.16, WT-CHLOR=15.17±0.75, CD-CHLOR=13.00±1.00]. As expected, body weight and brain weight of CD mice were significantly reduced when compared to their WT littermates, but these alterations were not alleviated by CHLOR chronic administration [Fig. 3-A and B; genotype effect, respectively: F(1,18)=46.44 and 102.6, p<0.0001; effects of treatment and of its interaction with genotype, all n.s.].

When brain analyses were performed, CD animals showed an altered morphology in the motor cortex and hippocampus, and CHLOR treatment did not normalize these alterations, neither the reduced amount of YFP+ expressing neurons (Fig. 4-A and B, qualitative evaluation), nor the reduced dendritic length in both areas [Fig.4-C and D, genotype effect in cortex and hippocampus, respectively: F(1,10)= 25.76 and 83.38, p<0.001].

**Fig. 4:**
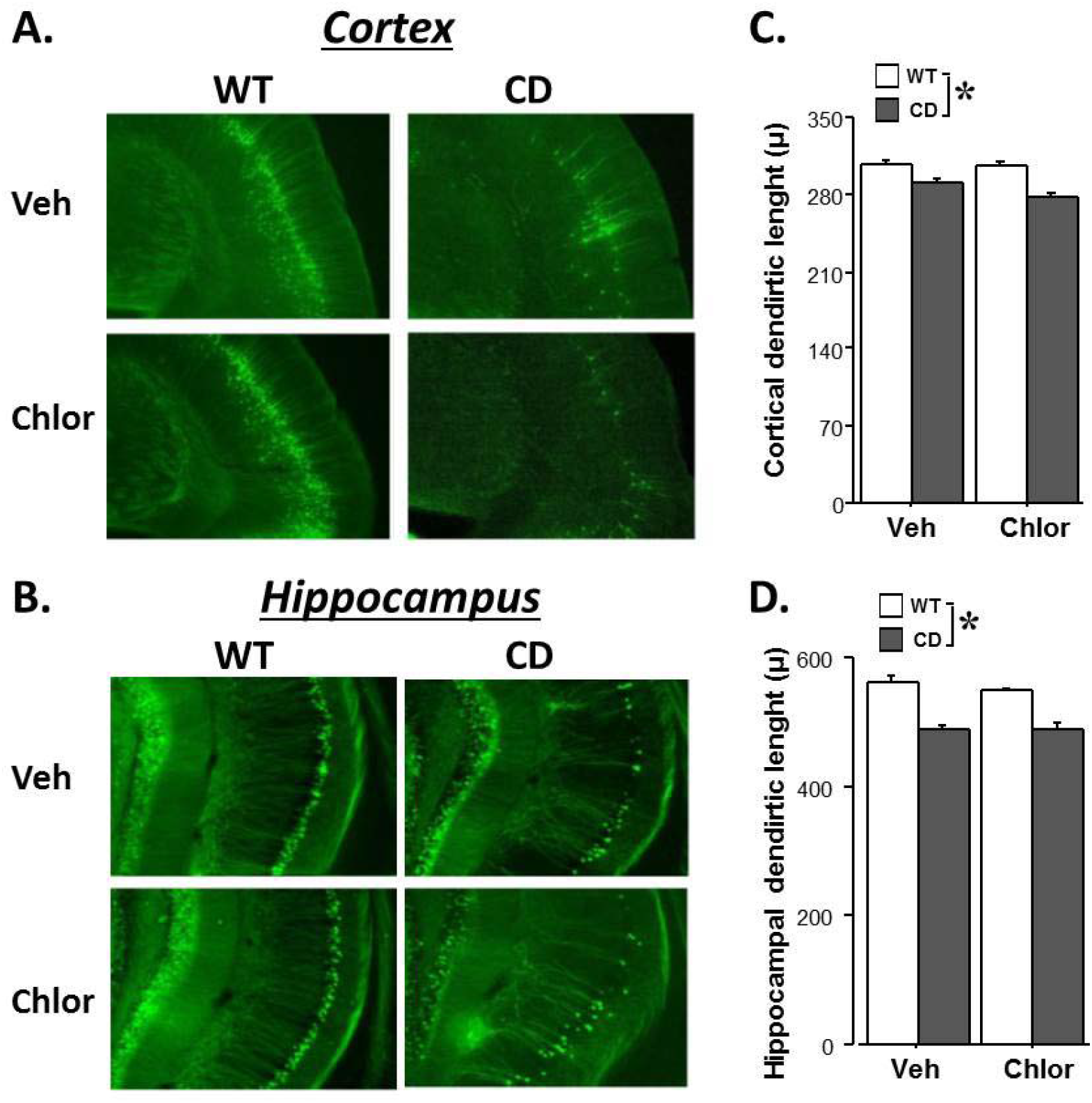
Chronic effects of chlorzoxazone (CHLOR; 5mg/Kg) on cortical and hippocampal morphology in male mice: Representative images of motor cortex (A; 4×) and hippocampus (B; 10×). Dendrite length of pyramidal neurons was measured in motor cortex (C) and in CA1 region (D). All measures were taken after 10 days of i.p. injections, with a 24hs interval from the last administration, n=3-4; *=p<0.05. Data are expressed as Mean±SEM.

**Fig. 5:**
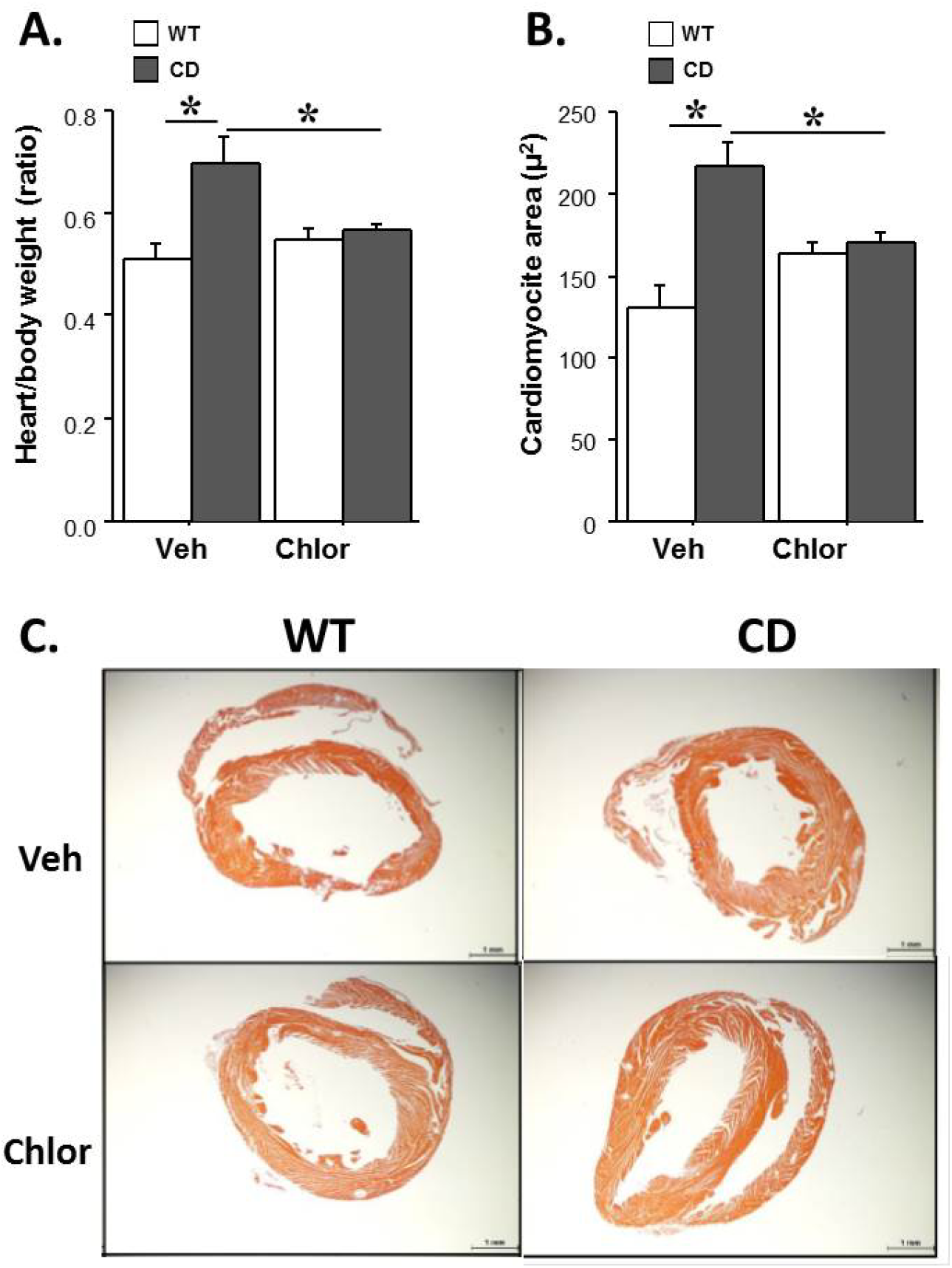
Chronic effects of chlorzoxazone (CHLOR; 5mg/Kg) on cardiac hypertrophy in male mice: Cardiac hypertrophy was quantified through assessment of heart/body weight ratios (A) and cardiomyocyte size (B). Representative images of hematoxylin-eosin stained transverse sections of heart samples reveal the thickening of the left ventricular wall in CD-VEH mice (C). Mice were sacrificed after 10 days of i.p. daily injections, with a 24hs interval from the last administration. n=4-6; *=p<0.05. Data are expressed as Mean±SEM.

CD mice were characterized by cardiac hypertrophy (Fig. 5), as demonstrated by the enhanced heart weight and the cardiomyocyte size compared to their WT littermates. These cardiac alterations were eliminated by chronic CHLOR administration, preventing both the enhanced heart weight [Fig. 5-A; genotype x treatment interaction: F(1,18)=7.50, p<0.05; post-hoc: CD-VEH versus WT-VEH and versus CD-CHLOR] and the enhanced cardiomyocytes’ area [Fig. 5-B; genotype x treatment interaction: F(1,15)= 14.19, p<0.01; post-hoc: CD-VEH versus WT-VEH and versus CD-CHLOR].

## DISCUSSION

Our data demonstrate for the first time the efficacy of Chlorzoxazone (CHLOR) in rescuing the behavioural and cardiac abnormalities of CD mice, i.e., the last generation mouse model of WBS. These results suggest that potassium and especially BKCa channels may be altered in CD mice, either in terms of the expression of channel subunits, as observed in *in vitro* studies on neuronal samples derived from WBS patients (Khattak, Brimble et al. 2015), and/or in terms of channel functionality, as observed in other NDDs (Laumonnier, Roger et al. 2006, Hebert, Pietropaolo et al. 2014, Zhang, Bonnan et al. 2014). Future studies are needed to specifically test this hypothesis, requiring electrophysiological and biochemical approaches to extensively assess potential alterations in the functional properties, as well as in the expression of BKCa channels in CD mice. These results will have to be then confirmed in WBS patients, in order to provide support to the hypothesis that channelopathies may be indeed a common etiopathological mechanism of multiple NDDs (Lee and Jan 2012). Our findings here provide convincing pre-clinical evidence for this hypothesis, as they fully match with the results we previously obtained in Fmr1-KO mice, showing the therapeutic effects of CHLOR (following the same experimental protocols used here) on behavioural FXS-like phenotypes (Lemaire-Mayo, Piquemal et al. 2020).

We described therapeutic effects of CHLOR on both male and female CD mice, although the rescue of cardiac hypertrophy appeared less marked in female mutants (as shown in the supplementary material section). Although most pre-clinical studies on NDDs focus on male mice only, and no difference between male and female CD mice has emerged so far in their neuro-cardio-behavioural phenotypes (Segura-Puimedon, Sahun et al. 2014), a more extensive evaluation of the effects of CHLOR in female CD mice would be of high relevance to fully assess potential sex differences in the response to therapies. This issue is of special interest since WBS, in contrast to other NDDs, does not preferentially or more markedly affect the male human population (Pankau, Partsch et al. 1992, Porter, Dodd et al. 2009), thus enhancing the importance of including female subjects in pre-clinical studies.

The therapeutic impact of chlorzoxazone on the behavioural phenotype of CD mice was demonstrated here both following acute and chronic administration, thus showing efficacy also outside the acute time window. Our data therefore suggest that long-term plasticity mechanisms may be triggered by the chronic administration of CHLOR. Brain-derived neurotrophic factor (BDNF) is a good candidate for future studies in this direction, as its reduced expression has been described in the hippocampal areas of CD mice, an alteration that has been already rescued together with behavioural abnormalities by other non-pharmacological therapeutic interventions (Ortiz-Romero, Borralleras et al. 2018). It is also possible that chronic CHLOR may eliminate the behavioural and cardiac alterations of CD mice by rescuing the altered calcium homeostasis described at the neuronal and cardiomyocyte levels in these mutants (Ortiz-Romero, Borralleras et al. 2018). Indeed, alterations in calcium homeostatic processes have been identified in WBS patients, including hyper-calcaemia and –calciuria (Sforzini, Milani et al. 2002), and have been recently suggested as an important player in the development of WBS pathological phenotypes (Ortiz-Romero, Borralleras et al. 2018).

While the identification of the specific mechanisms involved in the therapeutic effects of CHLOR requires further investigation, our data clearly demonstrate that the behavioural effects of this molecule are not mediated by a rescue of brain morphological alterations. CHLOR was indeed not able to attenuate the reduced brain weight or dendritic length found at both cortical and hippocampal levels in CD mice. As our treatment was limited to 10 days, starting at adult age, it is still possible that a rescue of these brain parameters (as well as of body weight loss) may require a longer period of administration and/or starting at earlier ages. Nonetheless, previous studies on therapeutic approaches in the CD model have also described behavioural and cardiac rescue with persisting alterations in brain weight and dendritic length/morphology (Ortiz-Romero, Borralleras et al. 2018). It is therefore possible that these brain morphological alterations may not play a crucial role in mediating the behavioural symptoms of WBS, or that their behavioural impact could be attenuated by activating alternative compensatory/plasticity brain mechanisms.

The ability of CHLOR to rescue both the behavioural and the cardiac alterations of CD mice, suggests that this molecule can induce effects not only at the central brain level. This is not surprising, considering that BKCa channels are known to control not only neuronal, but also muscular excitability and smooth muscle tone(Brenner, Perez et al. 2000). Nonetheless, this is the first evidence we obtained of non-behavioural effects of this molecule, as in our previous studies on Fmr1-KO mice we found therapeutic effects of CHLOR on the behavioural phenotypes, but not on other peripheral alterations, e.g., macro-orchidism (Lemaire-Mayo, Piquemal et al. 2020). Further investigations are needed to evaluate potential disease- or organ-specific effects of CHLOR, especially relative to the central versus peripheral activation of BKCa channels.

The therapeutic potential of CHLOR in treating both the cardiac and behavioural WBS-like symptoms is of crucial importance for the possible clinical applications of this drug. In fact, cardiovascular complications are the most common cause of health problems and even sudden death in WBS patients (Wessel, Gravenhorst et al. 2004). Moreover, the behavioral problems characterizing individuals with WBS can dramatically affect their quality of life, so that independent living is rarely achieved by these patients at adulthood (Morris, Demsey et al. 1988). Therefore, treatments that might alleviate the combination of these complex invalidating phenotypes deserve particular attention and converging research efforts.

## Supporting information

(as shown in the supplementary material section)

## FUNDING AND DISCLOSURE

This work has received funding from the Association “autour de Williams” 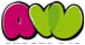 to SP and VC, and by Ministerio de Ciencia e Innovación (SAF2016-78508-R; AEI/MINEICO/FEDER, UE) to VC; MP, VLM, WC, and SP were funded also by CNRS and the University of Bordeaux. VLM, WC, EL, and SP are inventors on the patent: “Methods of treatment and/or prevention of disorders and symptoms related to BKCa and/or SK channelopathies” owned by Bordeaux University and assetsup.

## ACKNOWLEDGEMENTS

The authors thank Delphine Gonzales and the genotyping facility of Neurocentre Magendie, funded by Inserm and LabEX BRAIN ANR-10-LABEX-43, for animal genotyping. We thank Elodie Poinama and Renata Hermez for their expert animal care, Thierry Lafont for technical assistance, Christophe Halgand and Loic Grattier for informatics support.

## REFERENCES

Bai, Y. and X. Ma (2020). “Chlorzoxazone exhibits neuroprotection against Alzheimer’s disease by attenuating neuroinflammation and neurodegeneration in vitro and in vivo.” Int Immunopharmacol 88: 106790.

Bailey, C. S., H. J. Moldenhauer, S. M. Park, S. Keros and A. L. Meredith (2019). “KCNMA1-linked channelopathy.” J Gen Physiol 151(10): 1173–1189.

Belzung, C., S. Leman, P. Vourc’h and C. Andres (2005). “Rodent models for autism, a critical review.” Drug discovery today: Disease models 2(2): 93–101.

Borralleras, C., S. Mato, T. Amedee, C. Matute, C. Mulle, L. A. Perez-Jurado and V. Campuzano (2016). “Synaptic plasticity and spatial working memory are impaired in the CD mouse model of Williams-Beuren syndrome.” Mol Brain 9(1): 76.

Borralleras, C., I. Sahun, L. A. Perez-Jurado and V. Campuzano (2015). “Intracisternal Gtf2i Gene Therapy Ameliorates Deficits in Cognition and Synaptic Plasticity of a Mouse Model of Williams-Beuren Syndrome.” Mol Ther 23(11): 1691–1699.

Brenner, R., G. J. Perez, A. D. Bonev, D. M. Eckman, J. C. Kosek, S. W. Wiler, A. J. Patterson, M. T. Nelson and R. W. Aldrich (2000). “Vasoregulation by the beta1 subunit of the calcium-activated potassium channel.” Nature 407(6806): 870–876.

Carreno-Munoz, M. I., F. Martins, M. C. Medrano, E. Aloisi, S. Pietropaolo, C. Dechaud, E. Subashi, G. Bony, M. Ginger, A. Moujahid, A. Frick and X. Leinekugel (2018). “Potential Involvement of Impaired BKCa Channel Function in Sensory Defensiveness and Some Behavioral Disturbances Induced by Unfamiliar Environment in a Mouse Model of Fragile X Syndrome.” Neuropsychopharmacology 43(3): 492–502.

Dasilva, M., A. Navarro-Guzman, P. Ortiz-Romero, A. Camassa, A. Munoz-Cespedes, V. Campuzano and M. V. Sanchez-Vives (2020). “Altered Neocortical Dynamics in a Mouse Model of Williams-Beuren Syndrome.” Mol Neurobiol 57(2): 765–777.

Deng, L., H. Li, X. Su, Y. Zhang, H. Xu, L. Fan, J. Fan, Q. Han, X. Bai and R. C. Zhao (2020). “Chlorzoxazone, a small molecule drug, augments immunosuppressive capacity of mesenchymal stem cells via modulation of FOXO3 phosphorylation.” Cell Death Dis 11(3): 158.

Feng, G., R. H. Mellor, M. Bernstein, C. Keller-Peck, Q. T. Nguyen, M. Wallace, J. M. Nerbonne, J. W. Lichtman and J. R. Sanes (2000). “Imaging neuronal subsets in transgenic mice expressing multiple spectral variants of GFP.” Neuron 28(1): 41–51.

Gantois, I., A. Khoutorsky, J. Popic, A. Aguilar-Valles, E. Freemantle, R. Cao, V. Sharma, T. Pooters, A. Nagpal, A. Skalecka, V. T. Truong, S. Wiebe, I. A. Groves, S. M. Jafarnejad, C. Chapat, E. A. McCullagh, K. Gamache, K. Nader, J. C. Lacaille, C. G. Gkogkas and N. Sonenberg (2017). “Metformin ameliorates core deficits in a mouse model of fragile X syndrome.” Nat Med 23(6): 674–677.

Gaskill, B. N., A. Z. Karas, J. P. Garner and K. R. Pritchett-Corning (2013). “Nest building as an indicator of health and welfare in laboratory mice.” J Vis Exp (82): 51012.

Gaudissard, J., M. Ginger, M. Premoli, M. Memo, A. Frick and S. Pietropaolo (2017). “Behavioral abnormalities in the Fmr1-KO2 mouse model of fragile X syndrome: The relevance of early life phases.” Autism Res 10(10): 1584–1596.

Gribkoff, V. K., J. E. Starrett, Jr. and S. I. Dworetzky (2001). “Maxi-K potassium channels: form, function, and modulation of a class of endogenous regulators of intracellular calcium.” Neuroscientist 7(2): 166–177.

Halene, T. B., R. S. Ehrlichman, Y. Liang, E. P. Christian, G. J. Jonak, T. L. Gur, J. A. Blendy, H. C. Dow, E. S. Brodkin, F. Schneider, R. C. Gur and S. J. Siegel (2009). “Assessment of NMDA receptor NR1 subunit hypofunction in mice as a model for schizophrenia.” Genes Brain Behav 8(7): 661–675.

Hebert, B., S. Pietropaolo, S. Meme, B. Laudier, A. Laugeray, N. Doisne, A. Quartier, S. Lefeuvre, L. Got, D. Cahard, F. Laumonnier, W. E. Crusio, J. Pichon, A. Menuet, O. Perche and S. Briault (2014). “Rescue of fragile X syndrome phenotypes in Fmr1 KO mice by a BKCa channel opener molecule.” Orphanet J Rare Dis 9: 124.

Hohmann, N., A. Blank, J. Burhenne, Y. Suzuki, G. Mikus and W. E. Haefeli (2019). “Simultaneous phenotyping of CYP2E1 and CYP3A using oral chlorzoxazone and midazolam microdoses.” Br J Clin Pharmacol 85(10): 2310–2320.

Hu, H., L. R. Shao, S. Chavoshy, N. Gu, M. Trieb, R. Behrens, P. Laake, O. Pongs, H. G. Knaus, O. P. Ottersen and J. F. Storm (2001). “Presynaptic Ca2+-activated K+ channels in glutamatergic hippocampal terminals and their role in spike repolarization and regulation of transmitter release.” J Neurosci 21(24): 9585–9597.

Jackson, W. F. (2005). “Potassium channels in the peripheral microcirculation.” Microcirculation 12(1): 113–127.

Jimenez-Altayo, F., P. Ortiz-Romero, L. Puertas-Umbert, A. P. Dantas, B. Perez, E. Vila, P. D’Ocon and V. Campuzano (2020). “Stenosis coexists with compromised alpha1-adrenergic contractions in the ascending aorta of a mouse model of Williams-Beuren syndrome.” Sci Rep 10(1): 889.

Jirkof, P. (2014). “Burrowing and nest building behavior as indicators of well-being in mice.” J Neurosci Methods 234: 139–146.

Khattak, S., E. Brimble, W. Zhang, K. Zaslavsky, E. Strong, P. J. Ross, J. Hendry, S. Mital, M. W. Salter, L. R. Osborne and J. Ellis (2015). “Human induced pluripotent stem cell derived neurons as a model for Williams-Beuren syndrome.” Mol Brain 8(1): 77.

Kraeuter, A. K., P. C. Guest and Z. Sarnyai (2019). “The Nest Building Test in Mice for Assessment of General Well-Being.” Methods Mol Biol 1916: 87–91.

Latorre, R. and S. Brauchi (2006). “Large conductance Ca2+-activated K+ (BK) channel: activation by Ca2+ and voltage.” Biol Res 39(3): 385–401.

Laumonnier, F., S. Roger, P. Guerin, F. Molinari, R. M’Rad, D. Cahard, A. Belhadj, M. Halayem, A. M. Persico, M. Elia, V. Romano, S. Holbert, C. Andres, H. Chaabouni, L. Colleaux, J. Constant, J. Y. Le Guennec and S. Briault (2006). “Association of a functional deficit of the BKCa channel, a synaptic regulator of neuronal excitability, with autism and mental retardation.” Am J Psychiatry 163(9): 1622–1629.

Lee, H. Y. and L. Y. Jan (2012). “Fragile X syndrome: mechanistic insights and therapeutic avenues regarding the role of potassium channels.” Curr Opin Neurobiol 22(5): 887–894.

Lemaire-Mayo, V., M. Piquemal, W. E. Crusio, E. Louette and S. Pietropaolo (2020). “Therapeutic effects of Chlorzoxazone, a BKCa channel agonist, in a mouse model of Fragile X syndrome.” BiorRx 2020.12.11.389569.

Liu, Y. C., Y. K. Lo and S. N. Wu (2003). “Stimulatory effects of chlorzoxazone, a centrally acting muscle relaxant, on large conductance calcium-activated potassium channels in pituitary GH3 cells.” Brain Res 959(1): 86–97.

Martens, M. A., S. J. Wilson and D. C. Reutens (2008). “Research Review: Williams syndrome: a critical review of the cognitive, behavioral, and neuroanatomical phenotype.” J Child Psychol Psychiatry 49(6): 576–608.

Martindale, W. (1993). the Extra Pharmacopoeia. 30th Edition, Amer Pharmaceutical Assn.

Michalon, A., A. Bruns, C. Risterucci, M. Honer, T. M. Ballard, L. Ozmen, G. Jaeschke, J. G. Wettstein, M. von Kienlin, B. Kunnecke and L. Lindemann (2014). “Chronic metabotropic glutamate receptor 5 inhibition corrects local alterations of brain activity and improves cognitive performance in fragile X mice.” Biol Psychiatry 75(3): 189–197.

Mo, C., T. Renoir and A. J. Hannan (2015). “Novel ethological endophenotypes in a transgenic mouse model of Huntington’s disease.” Behav Brain Res 276: 17–27.

Moles, A. and R. D’Amato F (2000). “Ultrasonic vocalization by female mice in the presence of a conspecific carrying food cues.” Anim Behav 60(5): 689–694.

Morris, C. A., S. A. Demsey, C. O. Leonard, C. Dilts and B. L. Blackburn (1988). “Natural history of Williams syndrome: physical characteristics.” J Pediatr 113(2): 318–326.

N’Gouemo, P. (2014). “BKCa channel dysfunction in neurological diseases.” Front Physiol 5: 373.

Neely, C. L. C., K. A. Pedemonte, K. N. Boggs and J. M. Flinn (2019). “Nest Building Behavior as an Early Indicator of Behavioral Deficits in Mice.” J Vis Exp (152).

Oddi, D., E. Subashi, S. Middei, L. Bellocchio, V. Lemaire-Mayo, M. Guzman, W. E. Crusio, F. R. D’Amato and S. Pietropaolo (2015). “Early social enrichment rescues adult behavioral and brain abnormalities in a mouse model of fragile X syndrome.” Neuropsychopharmacology 40(5): 1113–1122.

Ortiz-Romero, P., C. Borralleras, M. Bosch-Morato, B. Guivernau, G. Albericio, F. J. Munoz, L. A. Perez-Jurado and V. Campuzano (2018). “Epigallocatechin-3-gallate improves cardiac hypertrophy and short-term memory deficits in a Williams-Beuren syndrome mouse model.” PLoS One 13(3): e0194476.

Pankau, R., C. J. Partsch, A. Gosch, H. C. Oppermann and A. Wessel (1992). “Statural growth in Williams-Beuren syndrome.” Eur J Pediatr 151(10): 751–755.

Pietropaolo, S., M. G. Goubran, C. Joffre, A. Aubert, V. Lemaire-Mayo, W. E. Crusio and S. Laye (2014). “Dietary supplementation of omega-3 fatty acids rescues fragile X phenotypes in Fmr1-Ko mice.” Psychoneuroendocrinology 49: 119–129.

Pietropaolo, S., A. Guilleminot, B. Martin, F. R. D’Amato and W. E. Crusio (2011). “Genetic-background modulation of core and variable autistic-like symptoms in FMR1 knock-out mice.” PLoS One 6(2): e17073.

Pober, B. R. (2010). “Williams-Beuren syndrome.” N Engl J Med 362(3): 239–252.

Porter, M. A., H. Dodd and D. Cairns (2009). “Psychopathological and behavior impairments in Williams-Beuren syndrome: the influence of gender, chronological age, and cognition.” Child Neuropsychol 15(4): 359–374.

Sah, P. and E. S. Faber (2002). “Channels underlying neuronal calcium-activated potassium currents.” Prog Neurobiol 66(5): 345–353.

Schubert, C. (2009). “The genomic basis of the Williams-Beuren syndrome.” Cell Mol Life Sci 66(7): 1178–1197.

Segura-Puimedon, M., I. Sahun, E. Velot, P. Dubus, C. Borralleras, A. J. Rodrigues, M. C. Valero, O. Valverde, N. Sousa, Y. Herault, M. Dierssen, L. A. Perez-Jurado and V. Campuzano (2014). “Heterozygous deletion of the Williams-Beuren syndrome critical interval in mice recapitulates most features of the human disorder.” Hum Mol Genet 23(24): 6481–6494.

Sforzini, C., D. Milani, E. Fossali, A. Barbato, G. Grumieri, M. G. Bianchetti and A. Selicorni (2002). “Renal tract ultrasonography and calcium homeostasis in Williams-Beuren syndrome.” Pediatr Nephrol 17(11): 899–902.

Tykocki, N. R., E. M. Boerman and W. F. Jackson (2017). “Smooth Muscle Ion Channels and Regulation of Vascular Tone in Resistance Arteries and Arterioles.” Compr Physiol 7(2): 485–581.

Wessel, A., V. Gravenhorst, R. Buchhorn, A. Gosch, C. J. Partsch and R. Pankau (2004). “Risk of sudden death in the Williams-Beuren syndrome.” Am J Med Genet A 127A(3): 234–237.

Zhang, Y., A. Bonnan, G. Bony, I. Ferezou, S. Pietropaolo, M. Ginger, N. Sans, J. Rossier, B. Oostra, G. LeMasson and A. Frick (2014). “Dendritic channelopathies contribute to neocortical and sensory hyperexcitability in Fmr1(-/γ) mice.” Nat Neurosci 17(12): 1701–1709.

