## Supplementary material for "Chlorzoxazone, A BKCa Channel Agonist, Rescues The Pathological Phenotypes Of Williams-Beuren Syndrome In A Preclinical Model": (as shown in the supplementary material section)

**SUPPLEMENTARY MATERIAL: THERAPEUTIC EFFECTS OF CHLORZOXAZONE IN FEMALE CD MICE**

Adult (4-5 months old) CD heterozygous females and their WT littermates were used as subjects, bred and maintained at the animal facilities of either Bordeaux (behavioural studies) or Barcelona (heart and brain analyses) Universities. All procedures, including breeding, drug administration, behavioural testing, heart and brain morphological assessment, were identical to those described for male studies in the main text. Estrous cycle was assessed on testing days by the analysis of vaginal smears, in order to exclude potential group (genotype x treatment) differences.

**Acute and chronic behavioural effects of CHLOR**

As expected and similarly to males, CD female mice treated with VEH showed deficits in motor abilities (Fig. S1-A and S2-A), reduced acoustic startle (Fig.S1-B) and enhanced anxiety levels (Fig.S2-B) and these behavioural abnormalities were treated by CHLOR, both at acute and chronic level. A single administration of CHLOR was able to markedly attenuate the motor deficits of CD mice in the rotarod test [interaction genotype x treatment: F(1,19)=9.56, p<0.01; post-hoc: CD-VEH versus WT-VEH and versus CD-CHLOR; Fig.S1-A], as well as their reduced sensory responsiveness at the highest stimulus levels [interaction genotype x treatment x stimulus intensity: F(3,150)=2.90, p<0.05; post-hoc at 90 dB: CD-VEH versus WT-VEH and versus CD-CHLOR; Fig.S2-B].

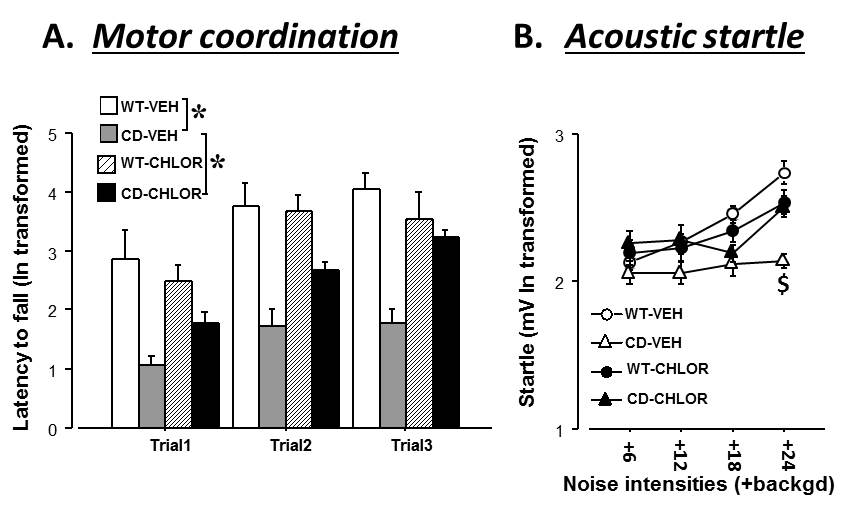

**Fig. S1**: Acute effects of chlorzoxazone (CHLOR; 5mg/Kg) on female behaviour: Behavioural tests were conducted 1h after a single i.p. injection and included rotarod test for motor coordination (A), and acoustic startle response test (B) with low intensity stimuli (background level=66dB). n=6-14; *=p<0.05. Data are expressed as Mean±SEM. $= versus CD-CHLOR and WT-VEH.

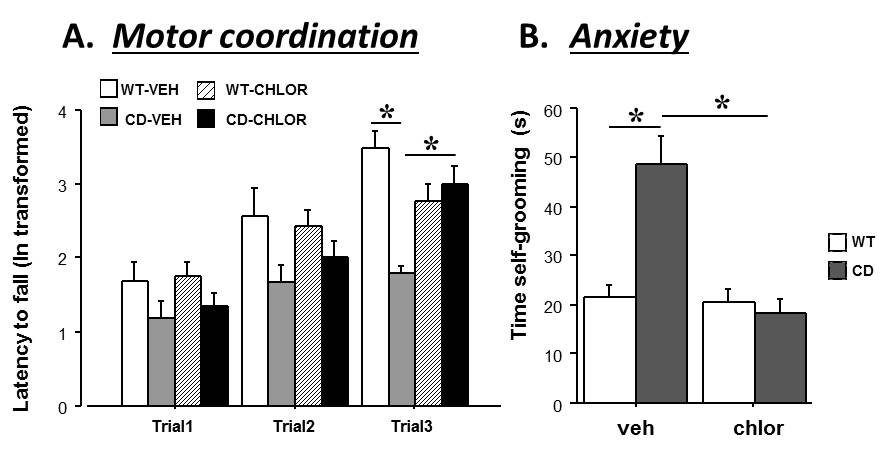

**Fig. S2**: Chronic effects of chlorzoxazone (CHLOR; 5mg/Kg) on behavioural deficits of female mice: Behavioural tests were conducted after 10 days of i.p. injections, with a 24hs interval from the last administration. They included the rotarod test for motor coordination (A), and anxiety measured by levels of self-grooming in a novel environment (B). n=8-10; *=p<0.05. Data are Mean±SEM.

**Chronic effects of CHLOR on body weight, brain and heart morphology**

As expected, body and brain weight of CD mice were significantly reduced when compared to their WT littermates, and these alterations were not alleviated by CHLOR chronic administration [Fig. S3-A and B; genotype effect, respectively, : F(1,24)=61.08 and 67.34, p<0.0001; effects of treatment and of its interaction with genotype, n.s., except for a significant interaction on body weight : F(1,24)=4.83, p<0.05, stemming exclusively on the slight reducing overall effect of CHLOR in WT mice]. CD mice were characterized by cardiac hypertrophy (Fig. S3-C and D), as demonstrated by the enhanced heart weight [genotype effect: F(1,22)= 11.34, p<0.01] and the cardiomyocyte size [genotype effect: F(1,19)= 38.63, p<0.01] compared to their WT littermates. This cardiac alteration was partially eliminated by chronic CHLOR administration, as it normalized the enhanced cardiomyocytes’ area [Fig. S3-D; genotype x treatment interaction: F(1,19)=14.69, p<0.01], but not the heart weight [Fig. S3-C; genotype x treatment interaction and treatment effect, all n.s.].

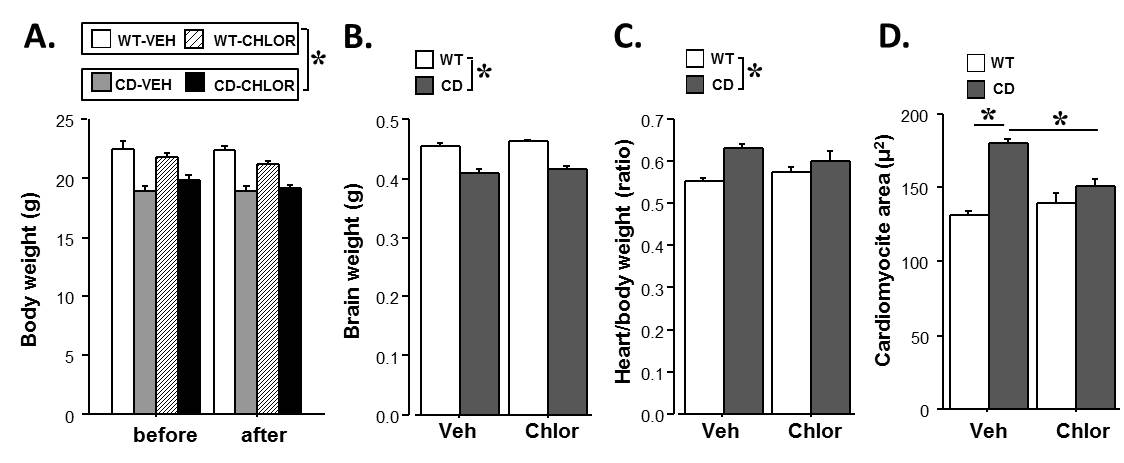

**Fig. S3**: Chronic anatomical effects of chlorzoxazone (CHLOR; 5mg/Kg): Measures of body (A), brain (B) and heart (C) weight, as well as of cardiomyiocyte size (D) were taken after 10 days of i.p. injections, with a 24hs interval from the last administration. n=5-9 for A, B and C; n=5-7 for D; *=p<0.05. Data are Mean±SEM.
